# Regenerating vascular mural cells in zebrafish fin blood vessels are not derived from pre-existing ones and differentially require *pdgfrb* signaling for their development

**DOI:** 10.1101/2021.03.27.437334

**Authors:** Elvin V. Leonard, Ricardo J. Figueroa, Jeroen Bussmann, Nathan D. Lawson, Julio D. Amigo, Arndt F. Siekmann

**Author notes:** Division Vascular Signaling and Cancer (A270), German Cancer Research Center, Heidelberg, 69120, Germany.

## Abstract

Vascular networks are comprised of endothelial cells and mural cells, which include pericytes and smooth muscle cells. It is well established that new endothelial cells are derived from pre-existing ones during the angiogenic phase of blood vessel growth. By contrast, mural cell ontogeny is less clear with an ongoing debate whether mural cells possess mesenchymal stem cell properties. To elucidate the mechanisms controlling mural cell recruitment during development and tissue regeneration, we studied the formation of zebrafish caudal fin arteries. Mural cells showed morphological heterogeneity: cells colonizing arteries proximal to the body wrapped around them, while those in more distal regions extended protrusions along the proximo-distal vascular axis. Despite these differences, both cell populations expressed platelet-derived growth factor receptor beta (Pdgfrb) and the smooth muscle cell marker myosin heavy chain 11a (Myh11a). Loss of Pdgfrb signalling during development or tissue regeneration resulted in a substantial decrease in mural cells at the vascular front, while those proximal to the body were less affected. Using lineage tracing, we demonstrate that precursor cells located in periarterial regions of the caudal fin and expressing Pgdfrb can give rise to mural cells, while in regeneration newly formed mural cells were not derived from pre-existing ones. Together, our findings reveal conserved roles for pdgfrb signalling in development and regeneration, while at the same time illustrating a limited capacity of mural cells to self-renew or contribute to other cell types during tissue regeneration.

## Introduction

During tissue regeneration, lost cell types need to be replaced through de-novo differentiation of tissue resident stem cells or from pre-existing differentiated cells (Poss, 2010). Several vertebrate species, such as zebrafish or newts, can regenerate a wide variety of tissues, such as appendages (Sehring and Weidinger, 2020; Tanaka, 2016), heart muscle (de Wit et al., 2020; Gonzalez-Rosa et al., 2017; Xiang and Kikuchi, 2016) or retina (Alunni and Bally-Cuif, 2016; Wan and Goldman, 2016). By contrast, mammals have lost most of their regenerative potential and repair injured tissues through imperfect wound healing, which can lead to scar formation and fibrosis (Eming et al., 2014; Erickson and Echeverri, 2018). During both processes blood vessel regrowth occurs, highlighting their high regenerative potential (Chavez et al., 2016; DiPietro, 2013, 2016). However, it is not clear whether the newly formed blood vessels are equally functional in both settings, and how they might influence tissue regeneration. Blood vessels consist of endothelial cells lining the inner vessel surface and ensheathing mural cells. Studies in the zebrafish fin (Kametani et al., 2015; Tu and Johnson, 2011; Xu et al., 2014) and in mouse arteries (McDonald et al., 2018) have shown that endothelial cells in regenerating blood vessels are derived from pre-existing ones. Similarly, during wound healing endothelial cells are being recruited from neighbouring blood vessels (Carmeliet, 2005; Tonnesen et al., 2000). These mechanisms closely resemble those observed during embryonic angiogenesis. By contrast, the ontogeny of newly forming murals cells and their contribution to blood vessel and tissue regeneration have remained unclear.

Mural cells can be subdivided into smooth muscle cells and pericytes, with intermediate cell types being present (Armulik et al., 2011; Grant et al., 2019; Holm et al., 2018). Generally, pericytes are important for guiding newly growing blood vessel sprouts during angiogenic phases (Eilken et al., 2017; Teichert et al., 2017) and later in blood vessel stabilization (Hellstrom et al., 2001). However, organ-depended functional differences exist. In the brain, they ensure blood-brain-barrier integrity (Langen et al., 2019) and regulate blood flow (Pfeiffer et al., 2021). Pericytes dying during an ischemic attack have been implicated in preventing tissue reperfusion (Hall et al., 2014). Renal pericytes contribute to ultrafiltration and blood pressure control, while hepatic stellate cells regulate sinusoidal blood flow and have immunoregulatory properties (Holm et al., 2018). Several of these functions have been attributed to distinct mural cell shapes with smooth muscle cells showing a perpendicular wrapping morphology in respect to the blood vessel axis, while pericytes align longitudinally with this axis (Armulik et al., 2011; Holm et al., 2018). Mural cells furthermore display heterogeneity in their origins, with those in the head region being neural crest derived (Sinha and Santoro, 2018). The mesothelium, the epithelium lining the coelomic cavity, generates most of the other mural cell populations with contributions from somitic tissue and the secondary heart field (Armulik et al., 2011). To which degree these differences in origin contribute to functional differences between mural cells remains to be determined. One master regulator of mural cell development is platelet-derived growth factor beta (pdgfb) signalling (Kazlauskas, 2017). Mural cells and their progenitors express Pdgfrb, while endothelial cells express pdgfb ligands (Gaengel et al., 2009). Loss of *pdgfb* or *pdgfrb* results in a reduction of pericyte numbers in a tissue specific manner without affecting their initial specification (Crosby et al., 1998; Hellstrom et al., 1999; Lindahl et al., 1997). Analysis of chimaeric animals suggests that smooth muscle cells similarly require *pdgfrb* signalling (Crosby et al., 1998). In zebrafish, mural cells can be detected along the trunk vasculature on the second day of development (Fortuna et al., 2015; Santoro et al., 2009), around 12 hours after their appearance on the cranial vasculature (Ando et al., 2016; Wang et al., 2014). Studies addressing the origins of mural cells showed that trunk and hindbrain pericytes are mesoderm-derived while those in more anterior regions have a neural crest origin (Ando et al., 2016; Stratman et al., 2017; Whitesell et al., 2014). Smooth muscle cells in zebrafish express similar marker genes when compared to mouse (Santoro et al., 2009), such as *tagln* (*SM22alpha*) (Li et al., 1996), *acta2* (Gabbiani et al., 1981) and *myh11a* (Miano et al., 1994). Zebrafish mural cells also require *pdgfrb* signalling during their development (see publication by Anjo et al. in this issue of *Development*). Together, these findings suggest that mural cell biology during developmental stages is well conserved in zebrafish.

Previous studies investigating mural cell function during tissue injury indicated that pericytes retained properties of mesenchymal stem cells with cultured pericytes being able to contribute an array of different cell types, such as chondrocytes (Crisan et al., 2008), adipocytes (Tang et al., 2008), skeletal muscle (Dellavalle et al., 2011; Dellavalle et al., 2007) and neurons (Dore-Duffy et al., 2006). More recent results, however showed that lineage-labelled pericytes did not contribute to these cell types in various injury settings in vivo, calling their stem cell properties into question (Guimaraes-Camboa et al., 2017). Despite these findings, work in zebrafish demonstrated that cells derived from *pdgfrb* expressing mural cells contribute a specific extracellular matrix promoting axonal growth after spinal cord injury (Tsata et al., 2021). Studies examining mural cells during wound healing suggest that differentiated pericytes are activated and proliferate, subsequently supporting angiogenesis, while at other times promoting fibrosis (Ansell and Izeta, 2015; Rodrigues et al., 2019). Thus, the functional significance of pericytes during tissue regeneration remains an area of debate. In addition, studies unequivocally providing evidence concerning the origin of newly forming pericytes either during wound healing or tissue regeneration have been lacking (Rodrigues et al., 2019). We also lack an understanding of the signalling pathways guiding mural cell development during regeneration and how impaired mural cell function might impact the function of newly formed blood vessels and tissues.

Here, we investigated mural cell morphology, ontogeny and the signalling pathways important for their development and regeneration in the zebrafish fin using newly developed transgenic lines. We show that zebrafish mural cells show two distinct morphologies depending on their proximo-distal location within the fin. They express the mammalian smooth muscle cell marker *myh11a* and can be lineage-traced to tissue resident progenitor cells. These express *pdgfrb*, but not *myh11a*. Further, mural cell differentiation during both fin development and regeneration relies on pdgfrb signalling. Lineage tracing experiments show that new perivascular cells in regenerating fins are not derived from pre-existing ones. We also do not detect significant contribution of differentiated mural cells to other cell types. Together, our studies illustrate the morphological diversity of mural cells in the zebrafish fin, while arguing against their stem cell properties. We further suggest that the signalling mechanisms patterning mural cells in juvenile stages are being redeployed during tissue regeneration.

## Results

### *Pdgfrb* expression marks cells with distinct morphologies and anatomical locations within zebrafish fins

During juvenile stages, the fin vasculature expands ventrally from an artery-vein loop in the posterior part of the embryo and continues to grow during adulthood (Figure 1a-e). To investigate mural cell development during these stages, we imaged double transgenic fish *Tg(Pdgfrb:citrine)*^*s1010*^; *(−0*.*8flt1:RFP)*^*hu5333*^. In these animals, arterial ECs express red fluorescent protein (RFP), while *pdgfrb* positive cells express citrine (Bussmann et al., 2010; Vanhollebeke et al., 2015). We observed the occurrence of citrine expressing cells along the artery as early as 1 week post fertilization (wpf; Fig. 1b, white arrowheads). These cells were present throughout juvenile stages at 2 wpf (Fig. 1c, c’, white arrowheads) and 3 wpf (Fig. 1d, d’, white arrowheads), as well as in 4 wpf fish (Fig. 1e-h’). In addition to mural cells, we detected distinct cell types with varying levels of citrine expression. One population consisted of ovoid cells either located at the fin ray base (Fig. 1c’, yellow arrowheads) or at the bone ray joints at later developmental time points (Figure 1g’’, yellow arrowheads). Moreover, we found cuboidal-shaped cells surrounding arterial blood vessels (Fig. 1f-h’, pseudocolored in light green in Fig. 1g’ and Fig. 1h’). In order to characterize and quantify these distinct cell populations, we divided the fin into 4 segments (S1-S4), starting with S1 closest to the fish body towards S4 at the distal end of the fin. We then measured cellular dimensions and distribution throughout these segments. This analysis showed that on average citrine expressing mural cells were longer and narrower than cuboidal-shaped cells (Fig. 1i). We also found that mural cell numbers increased from distal to more proximal locations (Fig. 1j). In order to more closely examine mural cells, we generated triple transgenic zebrafish (*Tg(pdgfrb:gal4)*^*ncv24tg*^; *(UAS:GFP)*^*nkuasgfp1a*^; *(−0*.*8flt:RFP)*^*hu5333*^). Analysis of these animals revealed additional morphological differences within the mural cell population. We found that GFP expressing cells either wrapped around arteries (Supplementary Fig. 1a-c’) or sent out long protrusions along the vessel axis (Supplementary Fig. 1d-e’, red arrowheads, see also Fig. 1g-h’). These cells were differentially distributed along the proximo-distal fin axis. While S4 was mostly devoid of mural cells, S3 contained mainly mural cells with long protrusions. Mural cells in segments closer to the body (S1, S2) were characterized by their wrapping morphology (Fig. 1k). To further characterize these morphologies, we made use of a double transgenic approach using *Tg(pdgfrb:gal4)*^*ncv24tg*^; *(UAS:KAEDE)*^*s1999t*^ fish. These fish express the photoconvertible protein Kaede in mural cell populations. Kaede photoconversion revealed complex cell shapes consisting of numerous finger-like extensions surrounding arteries (Fig. 1l, m). Wrapping mural cells either showed distinct borders between each other or overlapped (Fig. 1l, m). Thus, using citrine expression, as well as cell shapes as defining criteria, we were able to distinguish at least 4 different populations of *pdgfrb* expressing cells in developing zebrafish fins.

**Figure 1:**
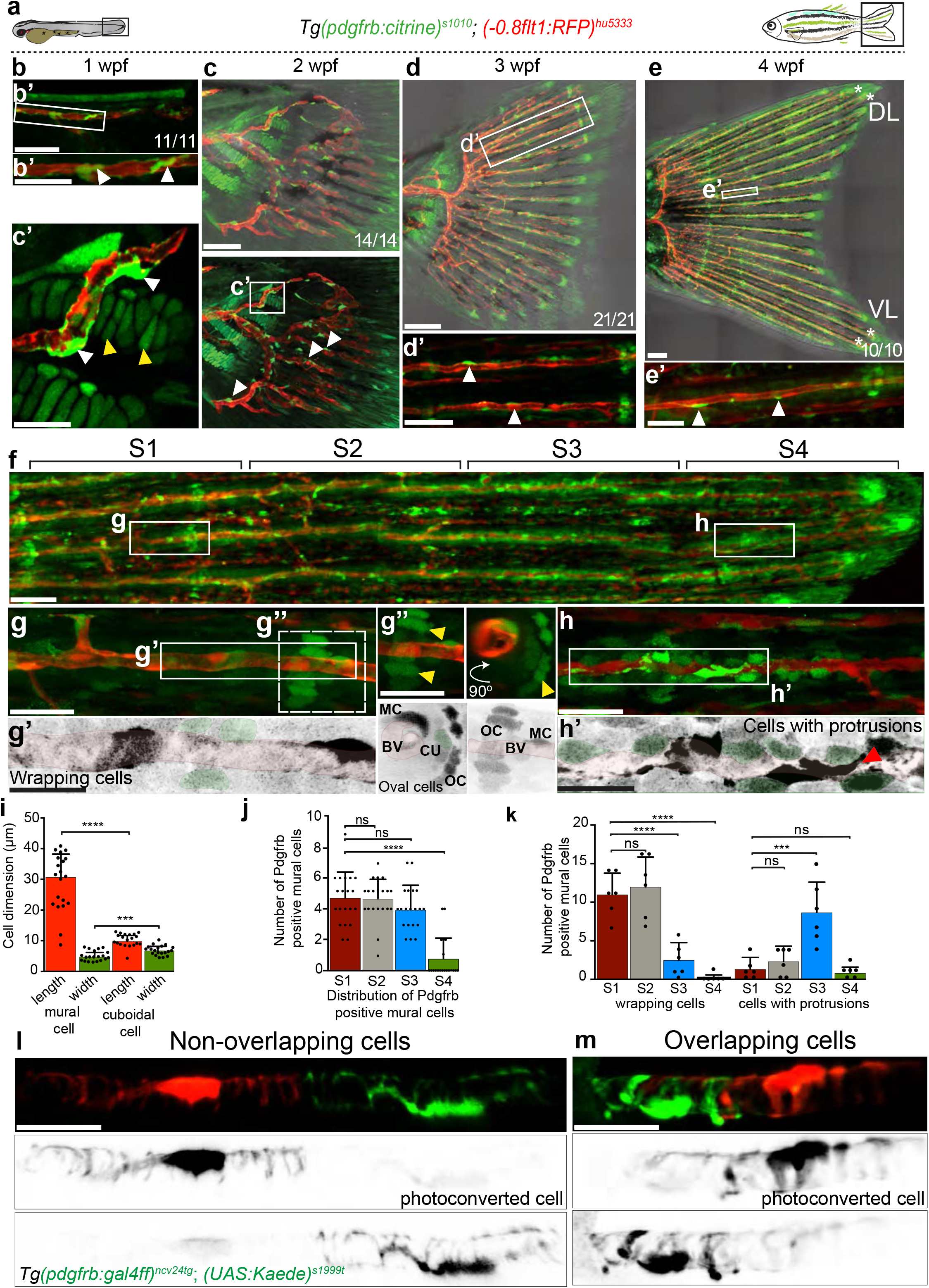
Pdgfrb expression marks distinct cell populations in the developing zebrafish fin. Maximum intensity projections of confocal z-stacks of *Tg (Pdgfrb:citrine)*^*s1010*^; *Tg(−0*.*8flt1:RFP)*^*hu5333*^ double transgenic fish labeling arterial ECs (red) and Pdgfrb positive cells (green) in lateral views with anterior to the left. (**a**) Schematic representation of the time course of the study. Mural cell recruitment to the developing caudal fin arteries was studied from 1 to 4 wpf. (**b**) Citrine expressing cells colonize axial vessels at 1 wpf. Scale bar: 20 µm. Boxed area enlarged in (**b’**) shows association of mural cells with arteries (white arrowheads). Scale bar: 10 µm. (**c**) Citrine expressing cells are recruited to the developing vasculature at 2 wpf. Scale bar: 30 µm. Boxed area enlarged in (**c’**) shows the association of individual mural cells with the artery (white arrowheads). Clusters of oval shaped cells formed at 2 wpf (yellow arrowheads). Scale bar: 10 µm. (**d**) Vasculature at 3 wpf. Scale bar: 70 µm. Dashed box region enlarged in (**d’**) shows the association of citrine expressing cells with arteries. Scale bar: 15 µm. (**e**) Caudal fin vasculature at 4 wpf, asterisks indicate the two blood vessels of the dorsal and ventral lobe used for quantifying citrine expressing cell distribution. Scale bar: 100 µm. Region enlarged in (**e’**) shows citrine expressing cells along arteries (white arrowheads). Scale bar: 30 µm. Numbers represent the individual embryos (or fish) analyzed for each developmental stage. (**f**) Caudal fin in 4 wpf fish. Scale bar: 20 µm. S1-S4 represent segments used for quantifying distribution of citrine expressing cells along blood vessels. (**g**) Proximal segment. Scale bar: 10 µm. Boxed region enlarged in (**g’**) shows citrine expressing cells wrapping around blood vessels. Scale bar: 7 µm. Dashed box region enlarged in (**g”**) shows citrine expressing cells (MC) on blood vessel (BV), oval shaped cells (OC) surrounding vessels (yellow arrowheads) and citrine expressing cuboidal shaped cells (CU, pseudo colored in green) between the mural cells and oval shaped cells. Scale bar: 5 µm. (**h**) Distal segment of caudal fin blood vessel. Scale bar: 10 µm. Boxed region enlarged in (**h’**) shows citrine expressing cells with protrusions elongating along the proximo-distal blood vessel axis (red arrowhead). Citrine expressing cuboidal cells are highlighted in green. Scale bar: 7 µm. (**i**) Classification of citrine expressing cells based on their dimensions. Citrine expressing mural cells are longer and narrower when compared to cuboidal cells. Mann-Whitney test, n=20 for each cell type. ***P=0.0001, ****P=<0.0001. (**j**) Quantification of citrine expressing mural cells across segments. One-way ANOVA n=20 caudal fin arteries from five different fish (average size: 1194 µm). ****P=<0.0001. (**k**) Quantifications of cell morphologies of *Tg(pdgfrb:gal4ff)*^*ncv24tg*^; *UAS(GFP)*^*nkuasgfp1a*^ expressing cells (See Supplementary Fig. 1). One-way ANOVA, n= 12 caudal fin arteries (two from dorsal or ventral lobe) from 6 individual fish (average size:1673µm) ****P=<0.0001, ***P=<0.0002. (**l, m**) Maximum intensity projections of confocal z-stacks of *Tg(pdgfrb:gal4ff)*^*ncv24tg*^; *(UAS:Kaede)*^*s1999t*^ mural cells. Cells wrapping around blood vessel are composed of non-overlapping (**l**) and overlapping cells (**m**).

### Co-expression of *pdgfrb* and *myh11a* defines mural cell populations

The classical perception of mural cell distribution along the vascular tree posits that proximal arterioles are covered by smooth muscle cells with their distinct morphologies surrounding arteries with their entire cell bodies, while more distally located capillaries are invested by pericytes (Grant et al., 2019; Holm et al., 2018). Based on this, we hypothesized that the mural cells we detected in distal segments of the fin most likely resembled pericytes, while we were unsure to which extend wrapping cells in more proximal locations would be similar to smooth muscle cells. To further investigate these identities, we developed a transgenic line expressing yellow fluorescent protein (YFP) under the control of the *myosin heavy chain 11a* (*myh11a*) promoter, a gene reported to be smooth muscle cell-specific (Miano et al., 1994; Vanlandewijck et al., 2018). Accordingly, when analyzing double transgenic *Tg(0*.*8flt1:RFP)*^*hu5333*^; *(myh11a:YFP)*^*mu125*^ zebrafish embryos at 5 days post fertilization (dpf), we detected YFP expression in mural cells surrounding arteries and in circulation (Supplementary Fig. 2a-c). Prominent YFP expression was also detectable in gut smooth muscle cells (Supplementary Fig. 2a, arrows). Similar to *Tg(Pdgfrb:citrine)*^*s1010*^ fish, we detected YFP expressing cells investing fin arteries as early as 1 wpf (Supplementary Fig. 2d-e’, yellow arrowheads) and throughout juvenile stages up to 4 wpf (Supplementary Fig. 2f-h’). We then quantified their morphology and distribution along the proximo-distal axis in 4 wpf larvae (Fig. 2a-f). This analysis revealed a striking similarity in respect to their distribution when compare to citrine expressing mural cells in *Tg(Pdgfrb:citrine)*^*s1010*^ fish. We detected both, wrapping cells (Fig. 2b, b’) and cells with protrusions (Fig. 2c, c’) expressing YFP. A greater number of cells displayed wrapping morphology in proximal regions, while more distal regions contained mostly cells exhibiting protrusions (Fig. 2d, d’). The most distal segment (S4) was again mostly devoid of mural cells (Fig. 2f). Thus, in the zebrafish fin, also mural cells without a characteristic smooth muscle cell morphology express YFP driven by the *myh11a* promoter.

**Figure 2:**
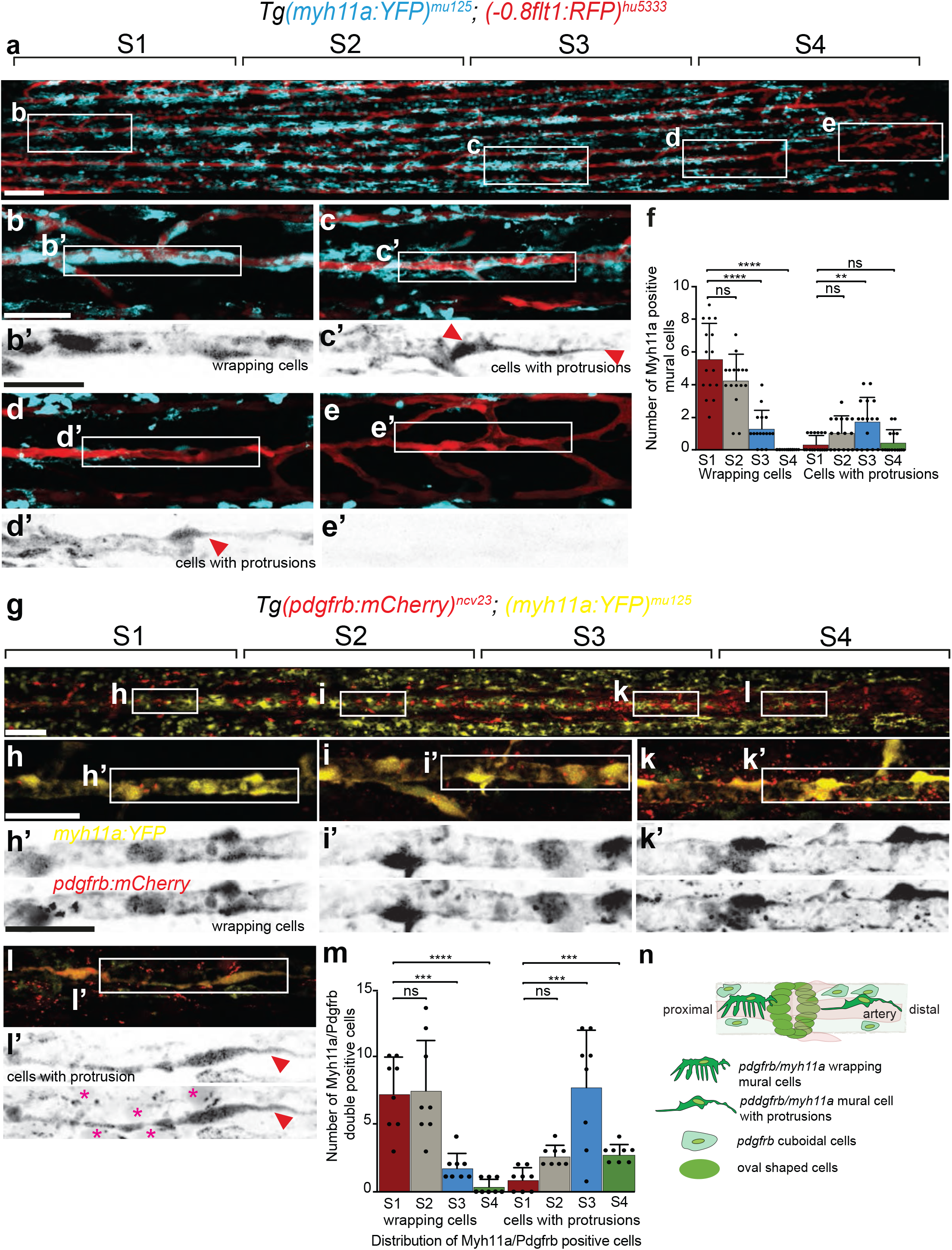
Colocalization of *myh11a* and *pdgfrb* expressing cells distinguishes vascular mural cells from other *pdgfrb* expressing cell populations. Maximum intensity projections of confocal z-stacks of *Tg(−0*.*8flt1:RFP)*^*hu5333*^; *Tg(myh11a:YFP)*^*mu125*^ double transgenic fish labeling arterial ECs (red) and YFP positive cells (blue) in lateral views with anterior to the left. (**a**) Caudal fin of 4 wpf fish. Scale bar: 100 µm. S1-S4 represent the four segments used for quantifying the distribution of YFP expressing cells. (**b**) Proximal segment of caudal fin blood vessel. Scale bar: 10 µm. Boxed region enlarged in (**b’**) shows YFP cells wrapping around blood vessel. Scale bar: 7 µm. (**c-e**) Mid vessel and distal segments of caudal fin blood vessel. Scale bar: 10 µm. Boxed region enlarged in (**c’-e’**) shows YFP positive cells with protrusions (red arrowheads) and absence of YFP expressing cells in most distal fin region. Scale bar: 7 µm. (**f**) Quantification of YFP positive cell distribution with wrapping and protrusion morphology. One-way ANOVA n=16 caudal fin arteries from 8 individual fish (average size: 1006 µm). ****P=<0.0001, **P=<0.0012, n.s., not significant, Error bars represent standard deviation (s.d.). (**g**) Maximum intensity projections of confocal z-stacks of *Tg (pdgfrb:mCherry)*^*ncv23*^; *Tg (myh11a:YFP)*^*mu125*^ double transgenic fish in lateral views with anterior to the left. Caudal fin of 4 wpf fish Scale bar: 100 µm. S1-S4 represent the four segments used for quantifying the distribution of *myh11a*/*pdgfrb* cells. (**h-l**) Proximal, mid vessel and distal segments of double transgenic fish. Scale bar: 10 µm. Boxed regions enlarged in (**h’-l’**) shows overlap of mCherry and YFP fluorophores in wrapping mural cells (**h’**-**k’**) and those extending protrusions (**l’**, red arrowhead). Note the presence of cuboidal cells that express only pdgfrb:mCherry (**I’**, magenta asterisks). Scale bar: 10 µm. (**m**) Quantification of the distribution of myh11a/pdgfrb positive cells with wrapping and protrusion morphology. One-way ANOVA, n=8 caudal fin arteries from 5 individual fish (average size:1938 µm). ****P=<0.0001, ***P=<0.000, n.s., not significant, Error bars represent standard deviation (s.d.). (**n**) Schematic illustrating the distribution of different fin cell populations based on their morphology and level of transgene expression.

To determine the relationship between *pdgfrb* and *myh11a* expressing cells, we generated double transgenic animals (*Tg(pdgfrb:mCherry)*^*ncv23*^; *(myh11a:YFP)*^*mu125*^), expressing mCherry under the control of the *pdgfrb* promoter and YFP in *myh11a* positive cells (Fig. 2g-l). All YFP expressing mural cells examined were positive for both transgenes. By contrast, YFP expressing cuboidal cells did not express mCherry (Fig. 2l’, magenta asterisks). We again detected a higher number of wrapping double positive cells in proximal fin regions, while distal fin regions mostly contained cells with protrusions (Fig. 2m). To determine whether *pdgfrb* and *myh11a* co-expression was a specific feature of fin mural cells, we analysed mural cells in the trunk of embryonic zebrafish at 6 dpf. In this setting, mural cells on arterial intersegmental vessels (aISV) co-expressed both transgenes, while those on venous ISVs only expressed the *pdgfrb* transgene (Supplementary Fig. 3). Together, these findings reveal that distinct mural cell morphologies in the zebrafish fin do not correlate with differences in the expression of the contractile protein Myh11a. Our findings further show that *pdgfrb* is more broadly expressed when compared to *myh11a* (Fig. 2n).

### *Pdgfrb* expressing cuboidal-shaped cells are precursors for vascular mural cells

We then set out to determine the cell population giving rise to fin mural cells. To do so, we generated transgenic fish that express the photoconvertible Dendra2 protein in the nuclei of *pdgfrb* positive cells (*Tg(Pdgfrb:H2B-dendra2)*^*mu158*^, Supplementary Fig. 4a-d’). Proof of concept studies in embryos showed that Dendra2 was expressed in mural cell populations in different embryonic regions (Supplementary Fig. 4b’-d’, white arrowheads) and could be photoconverted upon blue light exposure (Supplementary Fig. 4b’-d’, blue arrowheads). Based on their location close to arteries, we focused on cuboidal-shaped cells as potential mural cell precursors. In order to examine possible changes in cell shapes, we crossed *Tg(Pdgfrb:H2B dendra2)*^*mu158*^ fish with *Tg(pdgfrb:gal4ff)*^*ncv24tg*^; *(UAS:EGFP)*^*nkuasgfp1a*^ fish, expressing cytoplasmatic EGFP in *pdgfrb* positive cells. Examination of individual photoconverted cuboidal cells 5 days post photoconversion (dpc; Fig. 3) revealed that a proportion of photoconverted cells had undergone cell shape changes, now extending long processes along the vessel axis. In addition, we were able to detect proliferating cells, as evidenced by a doubling of red nuclei (Fig. 3c, c’). Thus, fin mural cells differentiate from local precursors that express *pdgfrb*.

**Figure 3:**
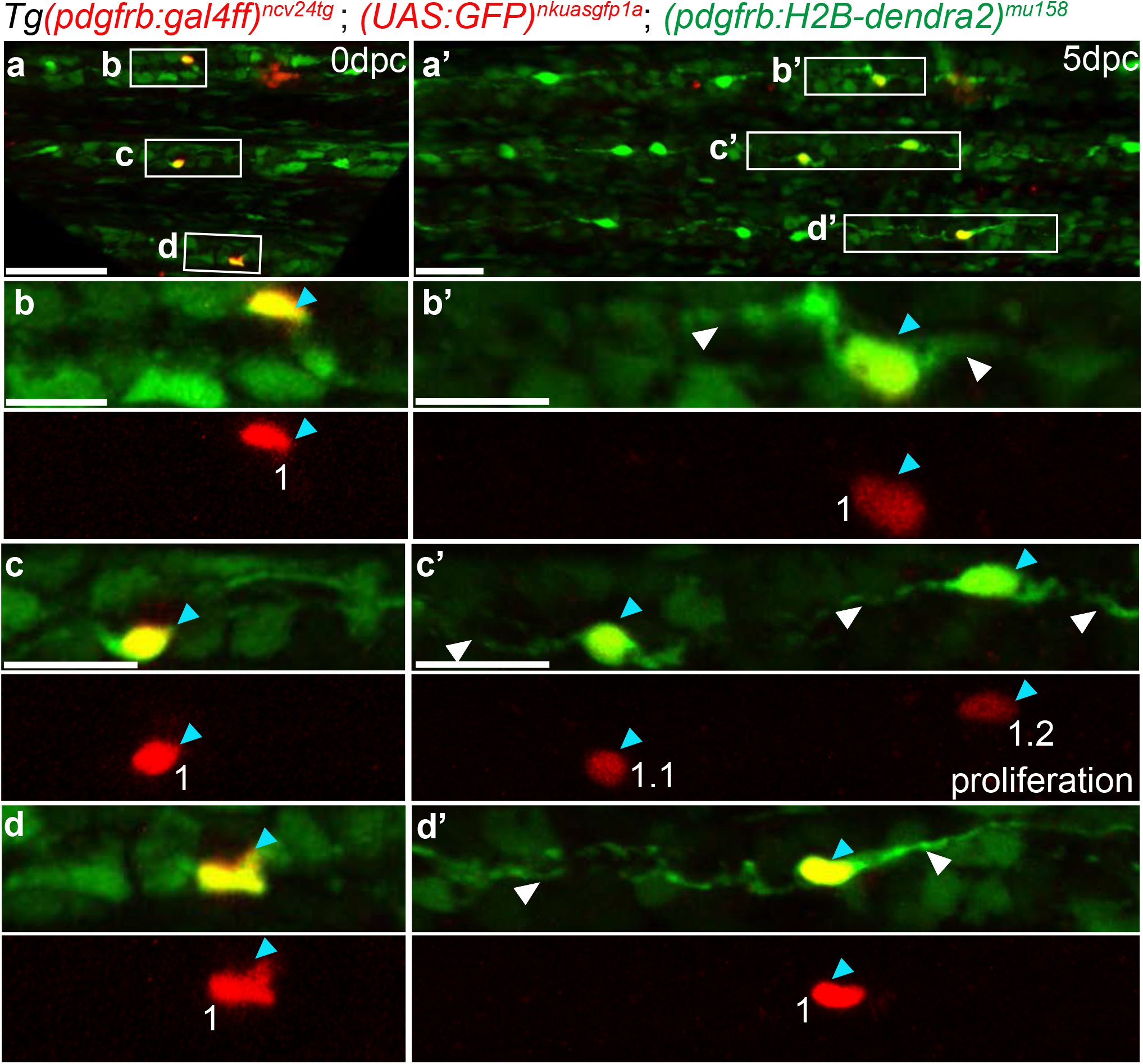
Cuboidal-shaped and pdgfrb expressing cells differentiate into mural cells during caudal fin artery development. Maximum intensity projections of confocal z-stacks of *Tg(pdgfrb:gal4ff)* ^*ncv24tg*^; *(UAS GFP)*^*nkuasgfp1a*^; *(pdgfrb:H2B-dendra2)*^*mu158*^ fins, labelling *pdgfrb* positive cells (green). (**a, a’**) Photoconverted nuclei at 0- and 5-days post conversion. Scale bar: 20 µm for (**a**), 30 µm for (**a’**). Boxed area enlarged in (**b-d**) shows photoconverted individual cells (blue arrowheads) at 0 dpc. Numbers mark individual cells. Scale bar: b: 15 µm (b), 20 µm (c, d). Boxed area enlarged in (**b’-d’**) shows photoconverted cells (blue arrowheads) extending protrusions (white arrowheads). Scale bar: 20 µm. (**c’**) Shows cell after division.

### Fin mural cell development in juvenile stages requires *pdgfrb* signaling

In order to determine the signaling pathways required for fin mural cell differentiation, we examined *pdgfrb*^*um148*^ mutant zebrafish (Kok et al., 2015). We divided fin arteries into four segments and quantified the distribution of mural cells using *Tg(Pdgfrb:citrine)*^*s1010*^; *(−0*.*8flt1:RFP)*^*hu5333*^ transgenic animals. In wildtype siblings, we observed a uniform distribution of mural cells along proximal and distal segments (Fig. 4a-c, yellow arrowheads), while cuboidal cells were present in distal segments (S3 and S4, Fig. 4d, blue arrowheads). By contrast, we observed a decrease in the number of mural cells colonizing distal blood vessel segments in *pdgfrb*^*um148*^ mutant fish, while mural cell numbers in proximal segments were unchanged (Fig. 4e, f, yellow arrowheads, quantified in Fig. 4i). In contrast, cuboidal cells along the blood vessel increased both in number and in expression levels of the citrine transgene (Fig. 4g,h, blue arrowheads, quantified in Fig. 4j). To corroborate our findings, we also analysed the distribution of mural cells in *Tg(−0*.*8flt1:RFP)*^*hu5333*^; *(myh11a:YFP)*^*mu125*^ transgenic fish. Similar to our observations for mural cells expressing *pdgfrb*, we detected a striking reduction in YFP expressing mural cells in distal segments of the fin in *pdgfrb* mutants (Fig. 4k-s). Therefore, also in the zebrafish fin, Pdgfrb signalling is instrumental for proper mural cell specification. However, proximal blood vessel segments were still invested with mural cells expressing transgenes driven by both *pdgfrb* and *myh11a* promoters. Our results further suggest a block in differentiation of cuboidal-shaped cells into the mural cell lineage in *pdgfrb* mutants, while at the same time leading to an upregulation of *pdgfrb* promoter activity in this precursor lineage.

**Figure 4:**
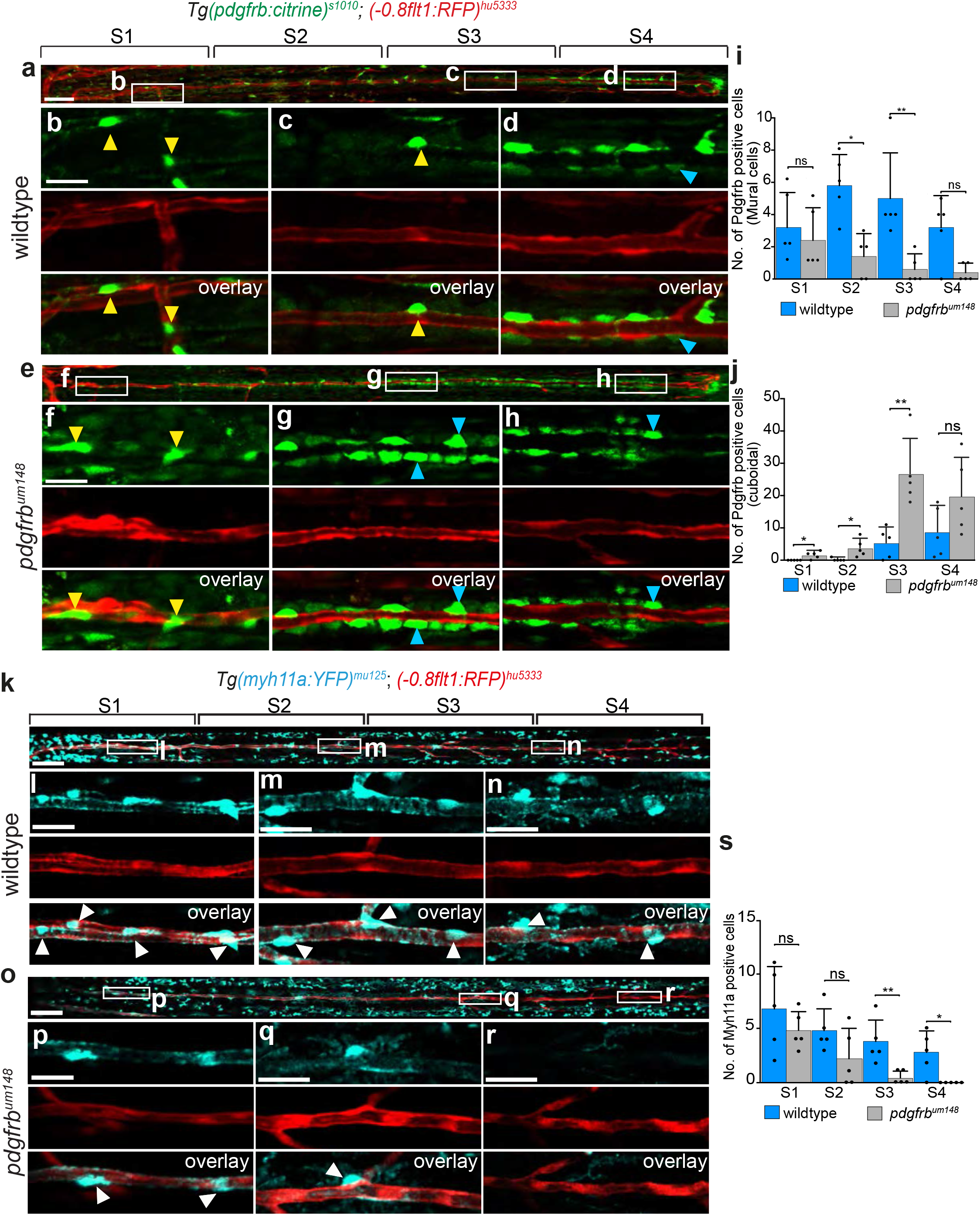
Mural cell recruitment to caudal fin arteries requires Pdgfrb signaling. Maximum intensity projections of confocal z-stacks of *Tg(−0*.*8flt1:RFP)*^*hu5333*^ ; *(pdgfrb:citrine)*^*s1010*^ fish. (**a**) Caudal fin artery in wildtype fish at 5 wpf. Scale bar: 70 µm. (**b, c**) Proximal and mid-vessel segments showing citrine expressing mural cells (yellow arrowheads). Scale bar: 10 µm. (**d**) Distal segment of caudal fin containing citrine expressing cuboidal cells (blue arrowhead). (**e**) Caudal fin vessel of *pdgfrb*^*um148*^ mutant fish at 5 wpf. Scale bar: 50 µm. (**f, g**) Proximal and mid-vessel segments contain citrine expressing mural cells (yellow arrowheads) and cuboidal cells (blue arrowheads). Scale bar: 10 µm. (**h**) Distal segment with cuboidal cells (blue arrowhead). (**i**) Quantification of Citrine expressing mural cell distribution across fin segments of wildtype and *pdgfrb*^*um148*^ fish. (**j**) Quantification of citrine expressing cuboidal cells in wildtype and *pdgfrb*^*um148*^ mutant fish. Mann-Whitney test, n=5 fish per group. n.s., not significant, Error bars represent s.d. *P<0.0476, **P=0.0079. (**k**) Maximum intensity projections of confocal z-stacks of *Tg(−0*.*8flt1:RFP)*^*hu5333*^; *(myh11a:YFP)*^*mu125*^ double transgenic fish. Scale bar: 80 µm. (**l, m**) Proximal and mid-vessel segments (boxed regions in (**k**)) shows YFP positive mural cells (white arrowheads). Scale bar: 15 µm (**k**), 10 µm (**l**). (**n**) YFP positive cells (white arrowheads) in distal segments. Scale bar: 10 µm (**o**) Caudal fin vessel of *pdgfrb*^*um148*^ mutants. Scale bar: 80 µm. (**p, q**) Proximal and mid-vessel segments showing YFP positive mural cells (white arrowheads). Scale bar: 15 µm (**o**), 10 µm (**p**). (**r**) Distal segment of caudal fin lacking YFP positive cells. Scale bar: 10 µm. (**s**) Quantification of YFP positive cells in wildtype and *pdgfrb*^*um148*^ mutant fish. Mann-Whitney test n=5 fish per group, n.s., not significant, error bars represent s.d. **P=0.0079, *P=0.0476.

### Regenerating fin mural cells require *pdgfrb* signaling

To examine whether the same signals contribute to mural cell development in tissue regeneration, we performed two types of fin regeneration experiments in wildtype and *pdgfrb* mutant animals. We either removed a piece of tissue containing a bone ray and an artery in the center of a fin ray (Fig. 5) or amputated about 50% of the fin (Supplementary Fig. 5). We then allowed the tissue to regenerate and determined mural cell numbers. In wildtype fish, we observed a newly forming artery as early as 3 days post injury (dpi; Fig. 5c), with a fully regenerated artery invested by mural cells being present at 5 dpi (Fig. 5d-f). When comparing wildtype fish (Fig. 5g-i, m) at 7 dpi to *pdgfrb*^*um148*^ mutants, we observed a significant reduction in mural cell recruitment to the artery in *pdgfrb* mutants (Fig. 5j-m). We also detected a reduction in mural cell coverage around regenerated arteries in the fin amputation assay (Supplementary Fig. 5). Thus, also in regenerative settings, *pdgfrb* signaling is a prerequisite for proper mural cell differentiation. These findings suggest that developmental signals are re-used during mural cell specification in regenerating tissues.

**Figure 5:**
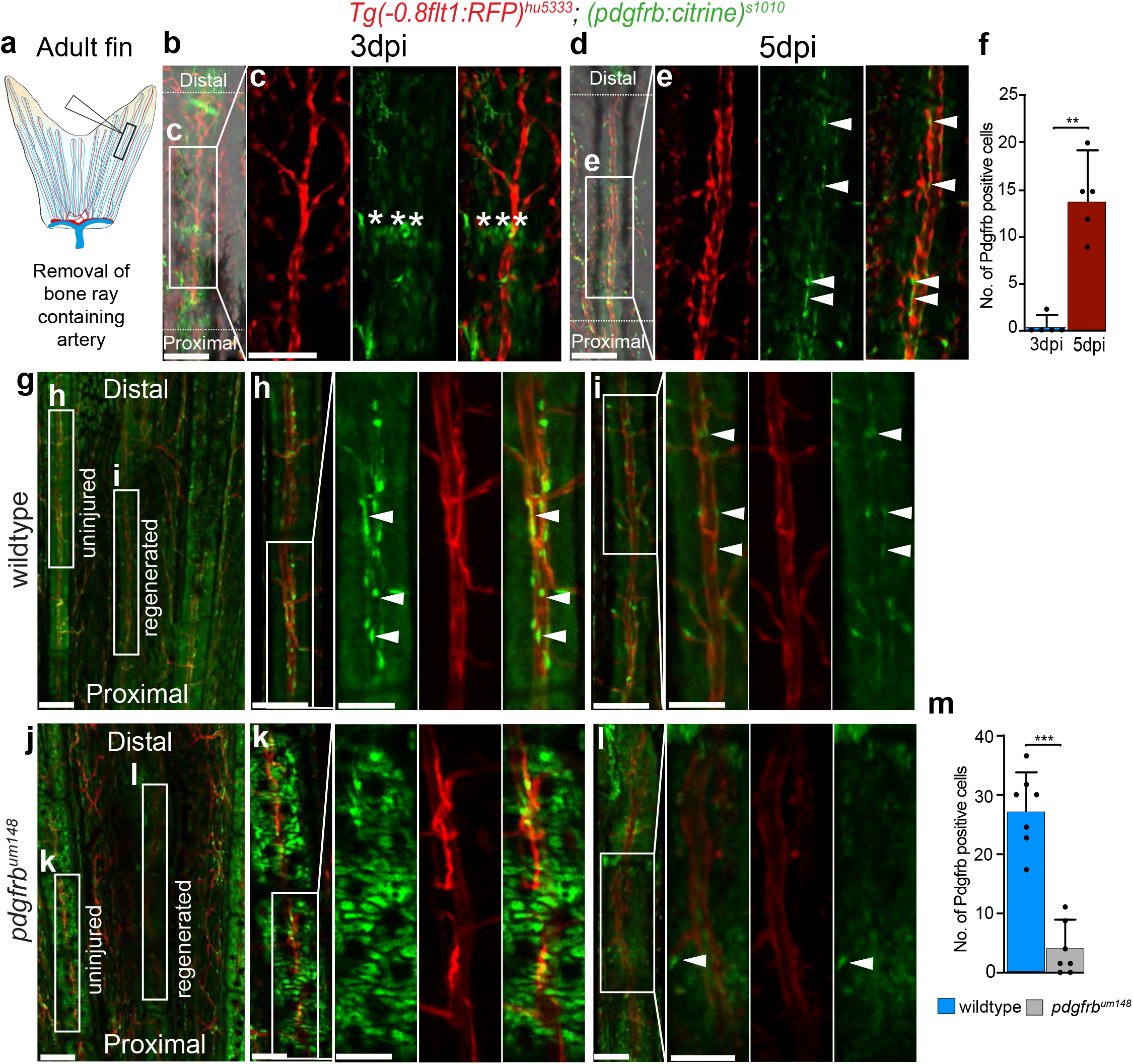
Pdgfrb signaling regulates mural cell recruitment during fin regeneration. **(a)** Schematic representation of the experimental approach to study blood vessel growth in regenerating tissue. Maximum intensity projections of confocal z-stacks of *Tg(−0*.*8flt1:RFP)*^*hu5333*^; *(pdgfrb:citrine)*^*s1010*^ fish. **(b)** Regenerating fin at 3dpi. Scale bar: 50 µm. (**c**) Citrine positive oval cells (asterisks) between bone ray segments. Scale bar: 30 µm. **(d, e)** Recruitment of mural cells (arrowheads) to the regenerating fin at 5 dpi. Scale bar: 50 µm. (**f**) Quantification of mural cells recruited to the regenerating blood vessels at 3 and 5dpi. Mann-Whitney test, n= 5 fish for each time point. n.s., not significant, error bars represent s.d. **P=0.0079. (**g**) Mural cell recruitment to in control wildtype fish. Scale bar: 50 µm. (**h**) Un-injured bone segment of wildtype fish. Scale bar: 20 µm. Boxed region enlarged shows the presence of mural cells (white arrowheads). Scale bar: 8 µm. (**i**) Regenerated bone segment at 7 dpi. Scale bar: 20 µm. Enlarged boxed region shows mural cells (white arrowheads). Scale bar: 8 µm. (**j**) Mural cell recruitment in *pdgfrb*^*um148*^ mutant fish. Scale bar: 50 µm (**k**) Un-injured bone segment. Scale bar: 20 µm. Enlarged boxed region shows cuboidal cells. Scale bar: 8 µm (**l**) Regenerated bone segment at 7 dpi. Scale bar: 20 µm. Enlarged boxed region shows decrease in mural cells (white arrowheads). Scale bar: 8 µm. (**m**) Quantification of the number mural in wildtype and *pdgfrb*^*um148*^ fish. Mann-Whitney test, n= 7 fish. Error bars represent s.d. ***P=0.0006

### Pre-existing mural cells are not precursors for regenerating mural cells

During regeneration, either tissue-resident stem stells or dedifferentiated lineage-restricted precursor cells are the source for new tissues and cell types (Tanaka and Reddien, 2011). To analyse whether new mural cells are derived from pre-existing ones, we tracked mural cells during regeneration through photoconverting dendra2 protein in *Tg(Pdgfrb:H2B-dendra2)*^*mu158*^ fish prior to injury (Fig. 6a-c). We then determined the contribution of photoconverted mural cells to the newly formed tissue at 5 dpi (Fig. 6d,e). We exclusively detected newly made green Dendra2 protein in the regenerated region of the fin, while the surrounding tissue still contained an abundance of previously photoconverted red cells (Fig. 6e). These findings indicate that pre-existing mural cells are not a major source of newly forming mural cells during tissue regeneration in the zebrafish fin (Summary of findings in Supplementary Fig. 6). They also suggest that these cells do not serve as stem cells during the regeneration of other tissues.

**Figure 6:**
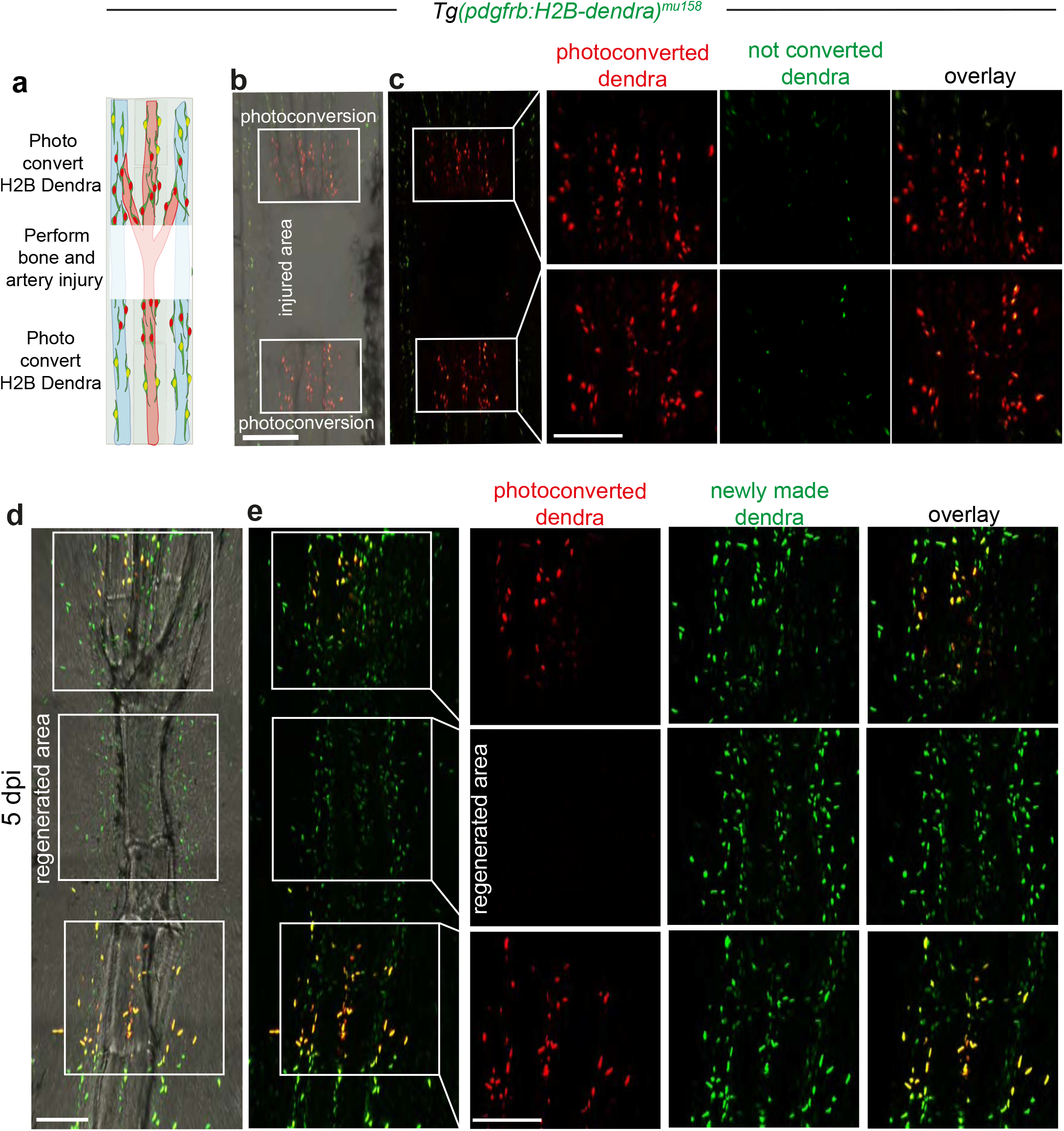
Pre-existing mural cells are not a major source of mural cells during fin regeneration. (**a**) Schematic representation of the experimental approach to study fin regeneration. Maximum intensity projections of confocal z-stacks of *Tg*(*pdgfrb:H2B-dendra2)*^*mu158*^ labelling *pdgfrb* expressing cells (green). (**b**) Fin ray after 12 hpi. Green to red photoconversion was performed above and below the site of injury. Scale bar: 100 µm (**c**) Photoconverted cells showing red dendra2 protein. Boxed regions enlarged shows the individual nuclei of photoconverted cells. Scale bar: 30 µm. (**d**) Regenerated fin ray at 5 dpi. Note the formation of a new bone ray segment. Scale bar: 50 µm (**e**) Newly made Dendra2 protein (green only) and previously photoconverted Dendra2 protein (green and red). Enlarged boxed regions show that previously photoconverted Dendra2 is restricted to regions proximal and distal to the site of tissue regeneration. Mural cells within the regenerated tissue are devoid of red Dendra2 protein. Scale bar: 30 µm

## Discussion

We have used newly generated transgenic zebrafish lines in order to investigate blood vessel mural cell populations in the zebrafish fin during adult stages and in tissue regeneration. We discovered the existence of two distinct mural cell populations that differed in respect to their morphology and location: in fin regions proximal to the fish body, we observed mural cells ensheathing arteries, while those in distal regions extended long processes along the proximo-distal vessel axis. These morphologies closely resemble those recently observed for subsets of pericytes on mammalian brain capillaries. Here, Grant et al. distinguish between thin-strand pericytes (TSP) sending out long processes and mesh pericytes (MP) that ensheath blood vessels (Grant et al., 2019). The authors furthermore distinguish two mural cell populations on upstream arterioles: ensheathing pericytes (EP) that ensheath blood vessels to a higher degree than MP and smooth muscle cells (SMC) displaying a characteristic morphology, encircling blood vessels. On note, we did not detect cells with a similar SMC morphology in the zebrafish fin. EPs and SMCs on arterioles express alpha smooth muscle actin, while TSPs and MPs on capillaries do not. While we did not include alpha smooth muscle actin in our analysis, we used *myh11a* as another marker previously reported to be exclusively expressed in differentiated smooth muscle cells (Miano et al., 1994). Expression domains of *Tg(myh11a:YFP)*^*mu125*^ are similar to those of alpha smooth muscle actin/acta2 in zebrafish embryos (Whitesell et al., 2014) and in the fin (Kametani et al., 2015), suggesting that there is overlap in the cell populations expressing contractile proteins. However, examining single cell sequencing data for mouse brain pericytes revealed that some also expressed *myh11*, while no *alpha smooth muscle actin/acta2* expression was detected in this cell population (Vanlandewijck et al., 2018). In line with this assessment, single cell sequencing of mural cell populations of different mouse organs has revealed a surprising diversification of pericytes, also in respect to the expression of components of the SMC contractile machinery, such as *myh11, tagln* and *acta2* (Muhl et al., 2020; Vanlandewijck et al., 2018). In organs such as the bladder or colon, pericytes expressed *tagln* and *acta2*, while in the brain and lung they did not. Therefore, the repertoire of contractile protein expression varies between pericytes and smooth muscle cell populations, as we now also show for mural cells in the fin of zebrafish. It will be of interest to evaluate the functional significance of these differences in the future.

Our results further suggest that fin mural cells might possess unique functions, possibly owing to the specific configuration of the fin vasculature. Fin blood vessels consist of arteries and veins that run through the entirety of the fin and are connected by intervessel commissures, but an elaborate capillary bed is missing, in particular in distal fin regions (Huang et al., 2003; Kametani et al., 2015; Xu et al., 2014). Further studies will be needed in order to decipher the functions of fin mural cells on vascular patterning and function and how they compare to other organs and tissues.

The finding that newly forming mural cells were not derived from pre-existing ones was surprising, since previous studies suggested that pericytes can function as mesenchymal stem cells, even though this notion was later called into question (Birbrair et al., 2017). During wound healing, several sources of pericytes, such as pre-existing pericytes or mesenchymal stem cells are being discussed (Morikawa et al., 2019). While a previous study showed that bone marrow-derived mesenchymal stem cells could contribute to the pericyte lineage during wound repair (Sasaki et al., 2008), definitive lineage tracing data concerning the contribution of pre-existing pericytes is still lacking. Our studies further did not find significant contribution of pre-existing mural cells to other cell types within regenerating fin tissue. This supports previous lineage tracing results in mice showing limited stem cell properties of mural cells (Guimaraes-Camboa et al., 2017). Thus, both in non-regenerative injury settings and during regeneration, differentiated mural cells do not seem to be a major source for newly forming tissues.

In keeping with the requirement for pdgfrb signalling during pericyte development in embryonic stages (Armulik et al., 2011), inhibition of pdgfrb signalling after wounding led to impaired pericyte recruitment (Rajkumar et al., 2006). These findings, together with our work showing deficient mural cell recruitment in *pdgfrb* mutants during tissue regeneration, underline the role of *pdgfrb* signalling as a master regulator of pericyte biology in different biological settings. Future studies will be necessary to illuminate possible differences and similarities in respect to the origins and functions of pericytes between wound healing and true tissue regeneration.

We found that mural cells in proximal fin regions still formed in *pdgfrb* mutants, while at the same time displaying a morphology distinct from perivascular cells in distal regions. This might suggest that distinct signalling pathways controlling mural cell recruitment and differentiation exist in different regions of the fin. These observations might further indicate separate embryonic origins for mural cells along the proximo-distal fin axis. Alternatively, distally located perivascular cells might give rise to proximal mural cell populations in wildtype settings, as suggested by a recent study investigating smooth muscle cell development in coronary arteries (Volz et al., 2015). Here, pericytes located on distal vessel branches are a precursor population for smooth muscle cells. In *pdgfrb* mutants, this route of distal to proximal differentiation would be compromised, leading to the activation of alternative sources for mural cells not normally used in wildtype fish. Detailed linage tracing studies in combination with single cell sequencing approaches will be needed in order to answer these questions and to determine how they relate to mural cells in other tissues and organisms.

## Acknowledgements

We would like to thank Naoki Mochizuki for sharing transgenic lines. We are grateful to Reinhild Bussmann, Mona Finch-Stephen, Bill Vought and Nadine Greer for excellent fish care.

## Contributions

E.V. L. and R.J.F. performed experiments. J.B. generated the *Tg(myh11a:YFP)*^*mu125*^ transgenic fish. N.D.L. generated *pdgfrb*^*um148*^ mutant fish. A.F.S. and J.A. conceptualized and supervised the work. E.V.L. and A.F.S. wrote the manuscript.

## Materials and Methods

### Zebrafish husbandry and strains

Zebrafish embryos were maintained in 1x E3 under standard husbandry conditions for 5 days in an incubator at 28.5 °C. After 5 days, the embryos were transferred to embryo tanks in the zebrafish facility, where they were raised until adulthood. Previously described zebrafish lines were *Tg(0*.*8flt1:RFP)*^*hu5333*^ (Bussmann et al., 2010), *Tg (pdgfrb:citrine)*^*s1010*^ (Vanhollebeke et al., 2015), *Tg (pdgfrb:mCherry)*^*ncv23*^, *Tg (pdgfrb:gal4ff)*^*ncv24*^ (Ando et al., 2016), *Tg(kdrl:TagBFP)*^*mu293*^ (Matsuoka et al., 2016), *pdgfrb*^*um148*^ (Kok et al., 2015). All animal experiments were performed in compliance with the relevant laws and institutional guidelines and were approved by local animal ethics committees of the Landesamt fur Natur, Umwelt und Verbraucherschutz Nordrhein-Westfalen.

### Generation of transgenic lines

Transgenic lines were generated using Bacterial artificial chromosome (BAC) recombineering. To generate the *Tg(myh11a:YFP)*^*mu125*^ and *Tg(Pdgfrb:H2B Dendra2)*^*mu158*^ the start codon of *myh11a* and *pdgfrb g*ene in BAC clone CH73-223E22 and CH1073-606I16 were replaced with YFP or H2B Dendra2 cassette using Red/ET recombineering (GeneBridges). The YFP cassette was amplified by PCR from pCS2+Citrine_kanR with the primers myh11a _HA1_citrine_fw (5′-ttatcaatcagtttttctccttaaatttgcagttgtcttaacccagcaccACCATGGTGAGCAAGGGCGAGGAG-3′) and myh11a_HA2_kanR_rev (5′-tctttgtccgtgaaaaggaatttctcatcatcgctcaagcctttcttcgtTCAGAAGAACTCGTCAAGAAGGCG-3′). iTol2_amp cassette was inserted into the vector back bone using a construct amplified with primers pTarBAC_iTol2_fw (5′-gcgtaagcggggcacatttcattacctctttctccgcacccgacatagatCCCTGCTCGAGCCGGGCCCAA GTG-3′) and pTarBAC_iTol2_rev (5′-gcggggcatgactattggcgcgccggatcgatccttaattaagtctactaATTATGATCCTCTAGATCAGAT C-3′) homology to the BAC vector is depicted in lowercase. The H2B Dendra2 cassette was amplified by PCR from pCS2+H2B Dendra2_kanR with primers Pdgfrb_HA1_dendra2_fw (5′-tgttgttttctctccgtctgcagtgttgaatgtgtcctgctctagaagaaCCACCATGCCAGAGCCAGCGAA) and Pdgfrb_HA1_dendra2_rev (5′-ttgtgatagcagtgaataggaagtggatgcggctgatggtcgaactcttTCAGAAGAACTCGTCAAGAAGG CG-3′) iTol2_amp cassette was inserted into the vector back bone using a construct amplified with primers pCC1FOS_ iTol2 _fw (5′-tctctgtttttgtccgtggaatgaacaatggaagtccgagctcatcgctaCCCTGCTCGAGCCGGGCCCAAGTG-3′) and pCC1FOS_ iTol2 _rev (5′-cgacacccgccaacacccgctgacgcgaaccccttgcggccgcatcgaatATTATGATCCTCTAGATCAG ATC). Homology to the BAC vector is depicted in lowercase. BAC DNA was purified by Midiprep (Invitrogen) on the day of injection a cocktail of 100 pg BAC DNA and 50 pg of tol2 transposase mRNA per embryo was injected into wild type embryos.

### Bone regeneration and fin amputation

Adult zebrafish were anaesthetized with in 0.02% tricaine and transferred to a petri plate. Bone resection was performed by removing a piece of bone ray enclosing an artery in the third or fourth fin ray of the dorsal or ventral lobe, using a scalpel. For fin amputation approximately 50% of the fin was amputated and the third fin ray from the dorsal or ventral lobe was used for analysis. The injured fish were kept in individual tanks until further experiments.

### Confocal microscopy

Live embryos or larvae (3-14dpf) were embedded in 1% low melting point agarose containing 0.02% tricaine. Amputated fins were embedded in 1% low melting point agarose. Fluorescent confocal images were acquired using a Zeiss LSM 780 /880 (objective lens: 20x Plan Apo NA 0.80 or 40x LD C Apochromat NA 1.10). For lineage tracing of cuboidal cells, we performed photoconversion in the distal regions of the juvenile fin. The anesthetized larvae were embedded in 0.75% agarose containing 0.02% tricaine. Individual cells were marked by regions and exposed to 8% of 405 nm laser excitation wavelength with 40 iterations, for 25 cycles. The larvae were removed from the glass plates and placed in fresh E3 medium and reimaged after 5 days. For lineage tracing of photoconverted Dendra2 cells during adult fin regeneration, bone resection was performed in adult fins and after 12 hours post injury anesthetized fish were transferred to glass plates and immobilized with 1.5% agarose. In order to keep fish sedated, E3 media containing 0.02% tricaine was added to the glass plate and the agarose around the gills, heart and mouth of the fish was carefully removed.

Photoconversion was performed by selecting regions around the resected bone with 10% of 405nm laser with 40 iterations, for 25 cycles. After photoconversion, fish were placed in individual tanks containing fresh E3. To speed up recovery, we used a pipette to transfer E3 to the mouth and gills of the fish. For photoconversion of Kaede expressing cells, amputated fin segments were embedded in agarose and photoconverted using a 405 nm laser with laser power of 8% and 30 iterations. Images for native or photoconverted Dendra2 and Kaede were acquired using laser settings for EGFP and RFP.

### Image processing and analysis

Acquired images were stitched using Zeiss ZEN software. Imaris software (Bitplane) was used to generate maximum intensity projections. The dimensions of cuboidal and mural cells with protrusions were determined by measuring their length and width. The distribution of mural cells that wrap or extend protrusions was quantified using Imaris software. Data were analyzed using Prism 9 (Graphpad) and graphs were plotted with mean standard deviation (SD). p values <0.05 were considered statistically significant. ImageJ (NIH) was used to generate monochrome images of different fluorophores. Adobe Illustrator software was used to compile images.

## Figure Legends

**Supplementary Figure 1:**
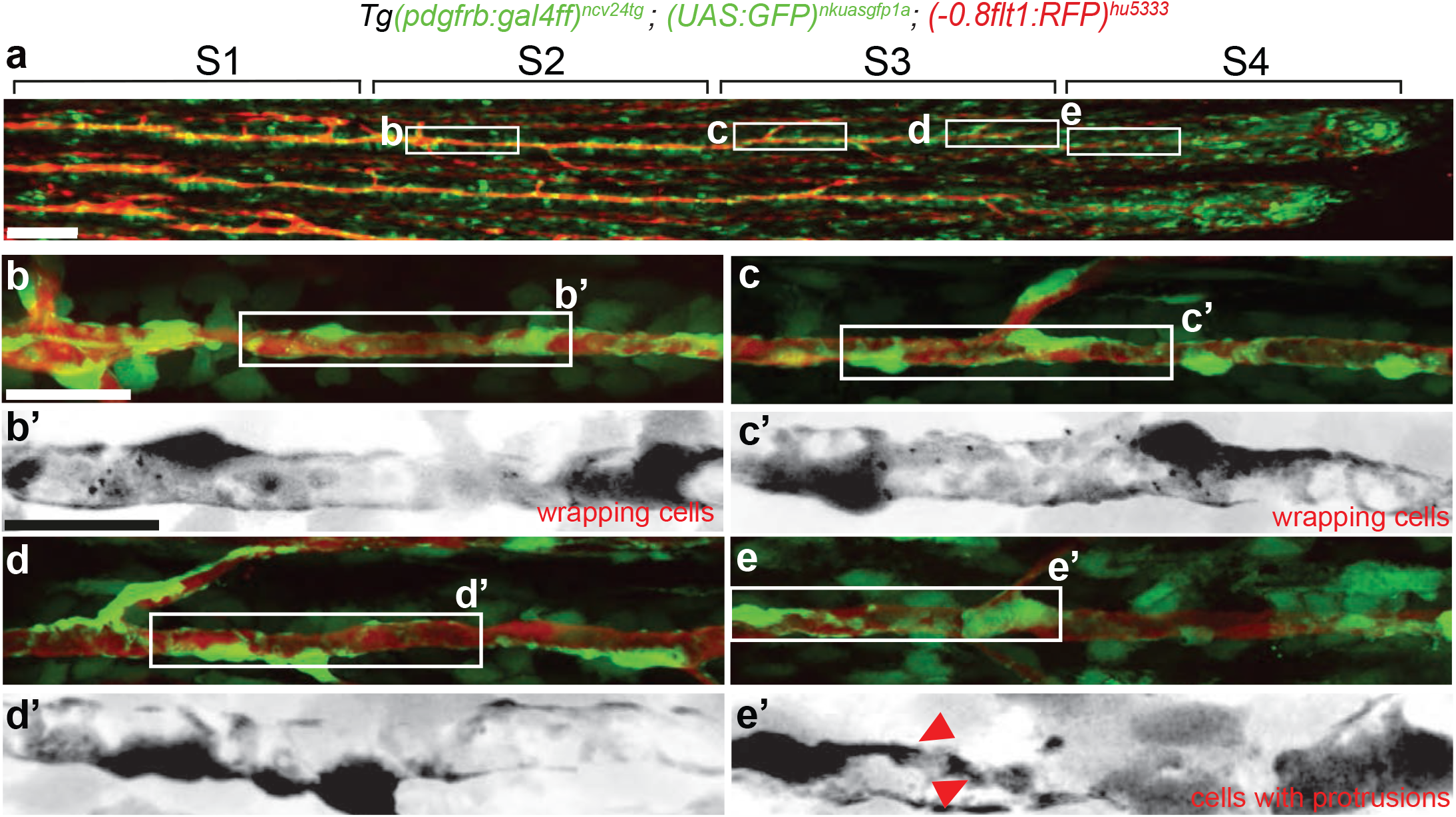
*Tg(pdgfrb:gal4ff)*^*ncv24tg*^; *(UAS:GFP)*^*nksuasgfp1a*^; *(−0*.*8flt1:RFP)*^*hu5333*^ transgenic fish mark mural cells on fin blood vessels. Maximum intensity projections of confocal z-stacks in lateral views with anterior to the left. (**a**) Caudal fin of 4 wpf fish. Scale bar: 100 µm. S1-S4 represent the four segments used for quantifying the distribution of GFP expressing cells. (**b**) Proximal segment of caudal fin blood vessel. Scale bar: 10 µm. Boxed region enlarged in (**b’**) shows GFP cells wrapping around blood vessel. Scale bar: 7 µm. (**c**-**e**) Mid vessel and distal segments of caudal fin blood vessel. Scale bar: 10 µm. Boxed regions enlarged in (**d’, e’**) show GFP positive cells with protrusions (red arrowheads). Scale bar: 7 µm. Quantification of images can be found in **Figure 1k**.

**Supplementary Figure 2:**
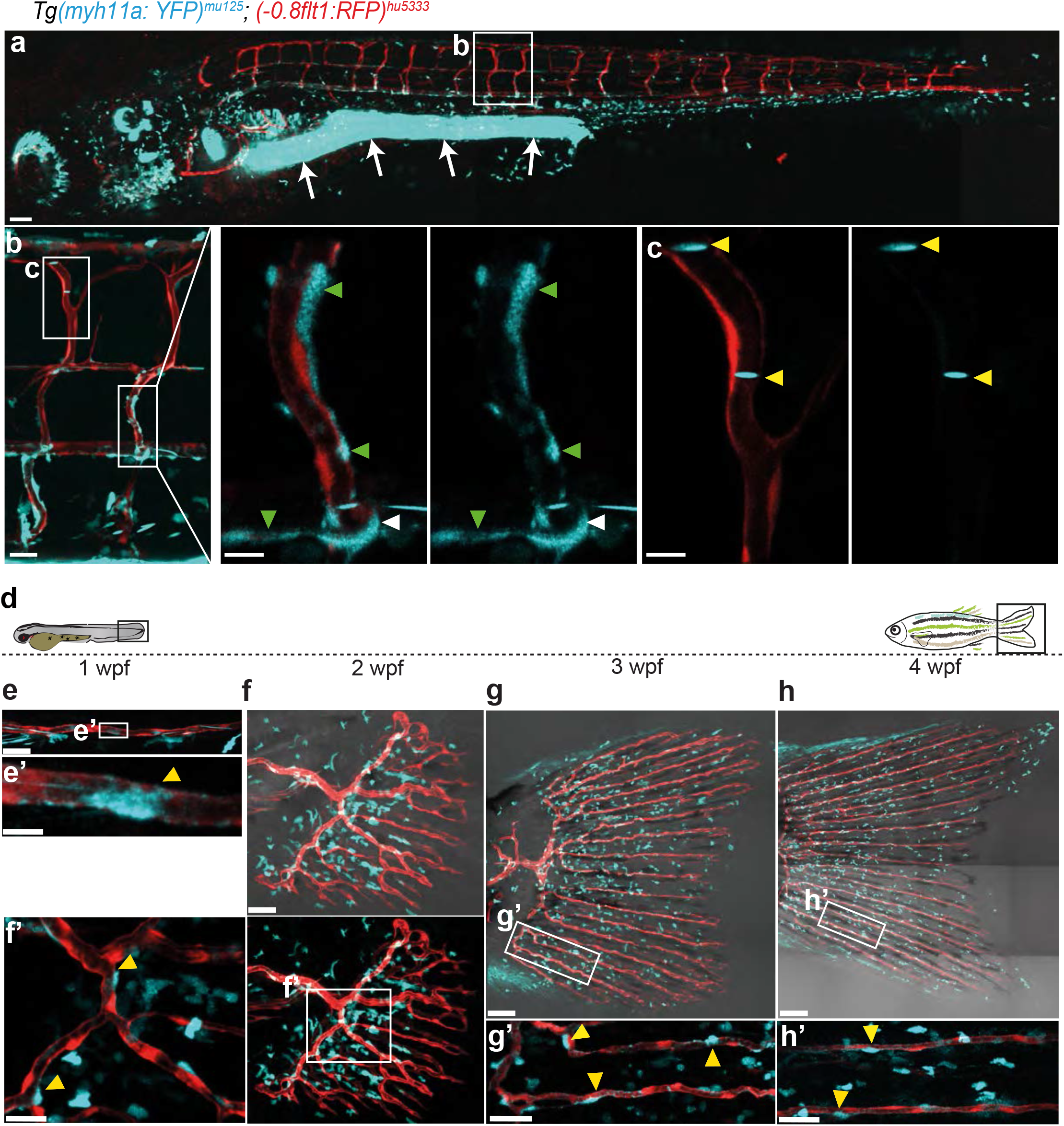
*Tg (myh11a:YFP)*^*mu125*^ marks mural cells of trunk and caudal fin vessels. Maximum intensity projections of confocal z-stacks of *Tg(−0*.*8flt1:RFP)*^*hu5333*^ ; *(myh11a:YFP)*^*mu125*^ double transgenic fish. (**a**) 5 dpf fish showing arterial ECs (red) and YFP positive cells (blue). Arrows mark expression in gut smooth muscle cells. Scale bar: 200 µm (**b**) Intersegmental vessels (ISVs). Scale bar: 20 µm. Enlarged boxed area shows YFP cells on arterial ISVs (green arrowheads) and on the ISV entrance from the dorsal aorta (white arrowhead). Scale bar: 5µm. (**c**) Expression of YFP in cells in the circulation (yellow arrowheads). Scale bar: 5µm. (**d**) Expression of YFP during juvenile fin development. Schematic representation of the time course of the study. (**e**) YFP positive cells colonize the caudal fin sprout by 1 wpf. Scale bar: 30 µm. Boxed area enlarged in (**e’**) shows the presence of YFP cells on arteries (yellow arrowheads). Scale bar: 8 µm. (**f**) Developing vasculature at 2 wpf. Scale bar: 30 µm. Boxed area enlarged in (**f’**) shows YFP positive cells colonizing blood vessels (yellow arrowheads). Scale bar: 8 µm. (**g**) Vasculature at 3 wpf. Scale bar: 50 µm. Boxed region enlarged in (**g’**) shows YFP in association with blood vessels (yellow arrowhead). Scale bar: 10 µm. (**h**) Caudal fin vasculature at 4 wpf. Scale bar: 100 µm. Boxed region enlarged in (**h’**) shows YFP cells on arteries (yellow arrowheads). Scale bar: 10 µm.

**Supplementary Figure 3:**
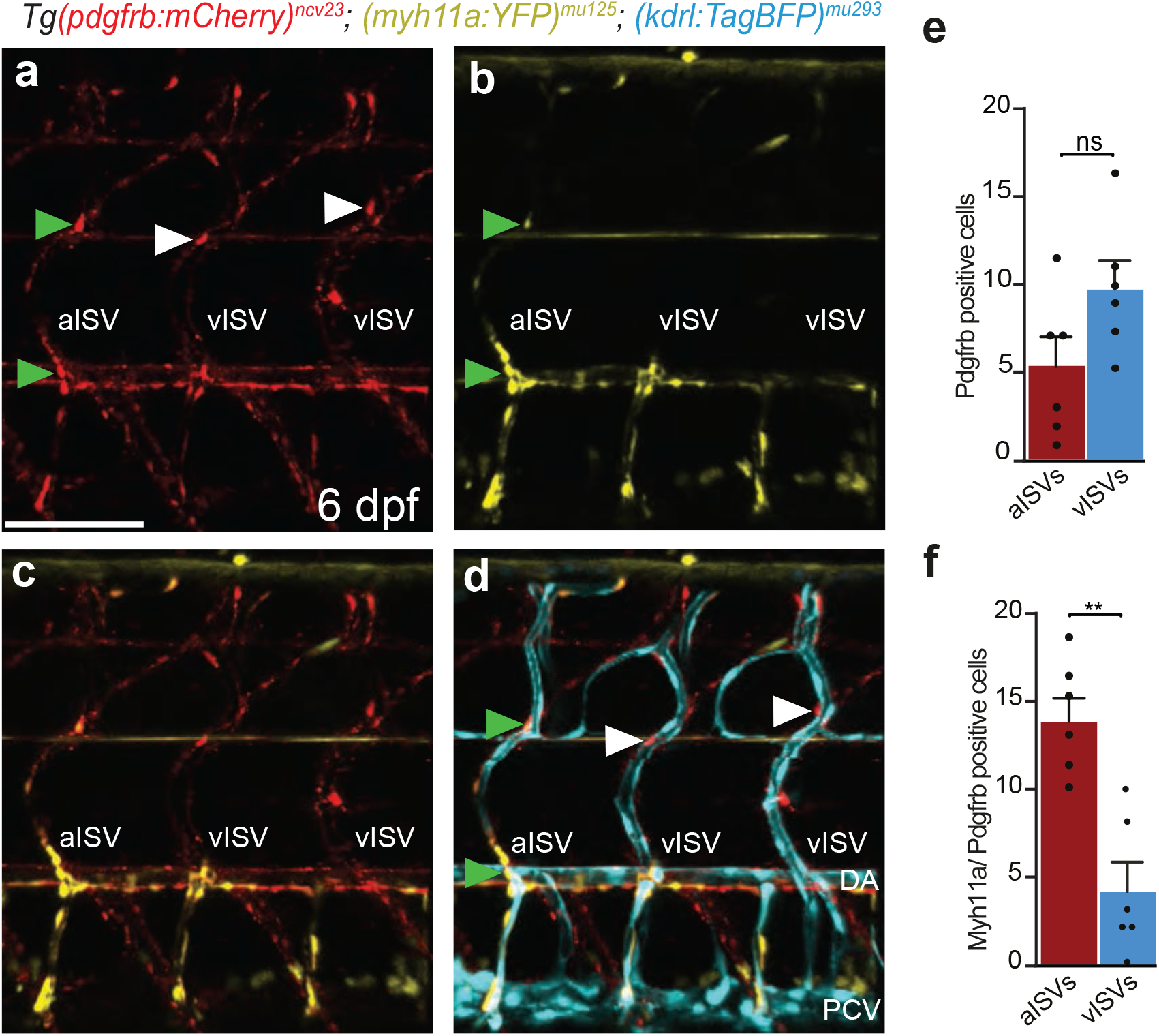
Colocalization of *myh11a* and *pdgfrb* in cells of embryonic trunk blood vessels. Maximum intensity projections of confocal z-stacks of *Tg(kdrl:TagBFP)*^*mu293*^ ; *(pdgfrb:mCherry)*^*ncv23*^; *(myh11a:YFP)*^*mu125*^ triple transgenic fish labeling all ECs (blue), *pdgfrb* positive cells (red) and *myh11a* positive cells (yellow). Lateral views, anterior to the left. (**a-d**) Trunk ISVs of 6 dpf zebrafish larvae show *pdgfrb* and *myh11a* positive cells associating with the vasculature. Scale bar: 50 µm. *pdgfrb*/*myh11a* double-positive cells associate with arterial intersegmental vessels (aISV) (green arrowheads). Myh11a expression is absent from mural cells associated with venous intersegmental vessels (vISV, white arrowheads). (**e**) Distribution of *pdgfrb* positive cells on ISVs. (**f**) Distribution of *myh11a*/*pdgfrb* double positive cells. Mann-Whitney test n=6. n.s., not significant, **P=0.0043, **P=0.0079, Error bars represent s.d.

**Supplementary Figure 4:**
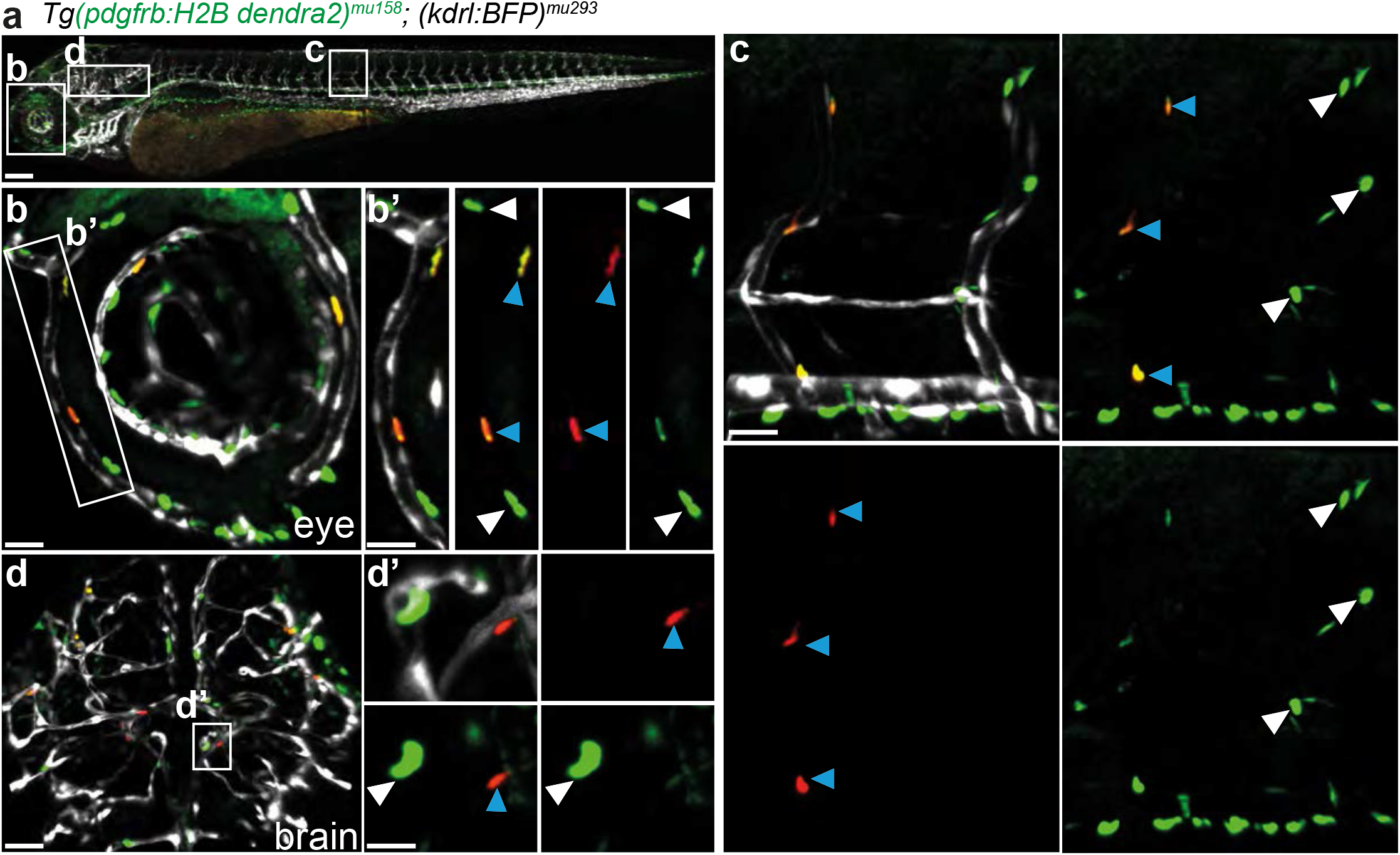
*Tg (pdgfrb:H2B dendra2)*^*mu158*^ labels cell nuclei of *pdgfrb* expressing cells with photoconvertible Dendra2 protein. Maximum intensity projections of confocal z-stacks of *Tg(pdgfrb:H2B dendra2)*^*mu158*^; *(kdrl:BFP)*^*mu293*^ double transgenic fish labeling the nuclei of pdgfrb positive cells with the photoconvertible protein Dendra2 (green) and all endothelial cells (blue). (**a**) 3 dpf embryo showing *pdgfrb* cells expressing Dendra2 protein in different vascular beds. Scale bar: 200 µm (**b**) Presence of Dendra2 positive cells along eye blood vessels. Scale bar: 20 µm. Boxed area enlarged in (**b’**) shows the inner optic circle with photoconverted cells (blue arrowheads) and unconverted cells (white arrowheads). Scale bar: 10 µm (**c**) Inter segmental vessels and dorsal aorta show un-converted (white arrowheads) and photoconverted (blue arrowheads) Dendra2 expressing cells. Scale bar: 15 µm (**d**) Head vessels of 48 hpf old zebrafish, anterior to top. Scale bar: 20 µm. Boxed regions enlarged in (**d’**) shows un-converted (white arrowheads) and photoconverted (blue arrowheads) cells on the central arteries. Scale bar: 10 µm.

**Supplementary Figure 5:**
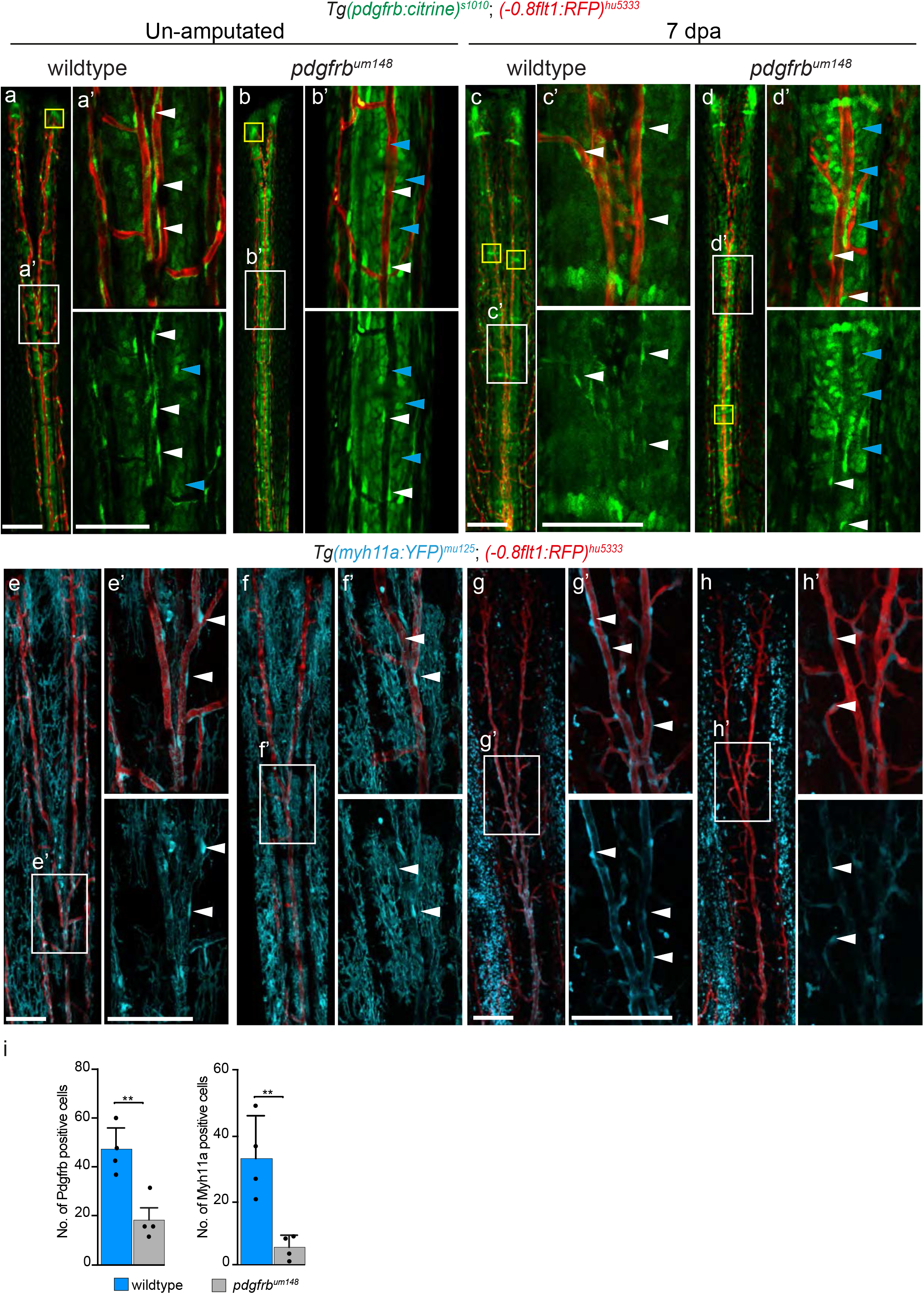
Mural cell recruitment to un-amputated and regenerated fin rays of the caudal fin. Maximum intensity projections of confocal z-stacks of *Tg(pdgfrb:citrine)*^*s1010*^; *(−0*.*8flt1:RFP)*^*hu5333*^ double transgenic fish labeling arterial ECs (red) and *pdgfrb* positive cells (green). (**a, b**) Caudal fin blood vessels of wildtype and *pdgfrb*^*um148*^ mutant fish. Scale bar: 50 µm. Note the presence of oval-shaped cells in terminal regions of the fin ray (yellow box). Boxed regions enlarged in (**a’** and **b’**) show presence of mural cells along blood vessels (white arrowheads). Cuboidal cell numbers are increased in mutants (blue arrowheads). Scale bar: 40 µm. (**c, d**) Regenerated caudal fin blood vessels of wildtype and *pdgfrb*^*um148*^ mutants at 7 days post amputation (dpa). Scale bar: 50 µm. Boxed regions enlarged in (**c’** and **d’**) show presence of mural cells along regenerated blood vessels (white arrowheads). Scale bar: 40 µm. Note increase of cuboidal cells in mutants (blue arrowheads). Maximum intensity projections of confocal z-stacks of *Tg(−0*.*8flt1:RFP)*^*hu5333*^ ; *(myh11a:YFP)*^*mu125*^ double transgenic fish labeling arterial ECs (red) and *myh11a* positive cells (blue). (**e, f**) Caudal fin blood vessels of wildtype and *pdgfrb*^*um148*^ mutants. Scale bar: 150 µm. Boxed regions enlarged in (**e’** and **f’**) show the presence of mural cells along blood vessels (white arrowheads). Scale bar: 40 µm. (**g, h**) Regenerated caudal fin blood vessels of wildtype and *pdgfrb*^*um148*^ mutants at 7 dpa. Scale bar: 500 µm. Boxed regions enlarged in (**g’** and **h’**) show mural cells (white arrowheads). Scale bar: 50 µm. (**i**) Quantification of *pdgfrb* and *myh11a* positive cells recruited to regenerated blood vessels. **P<0.005.

**Supplementary Figure 6:**
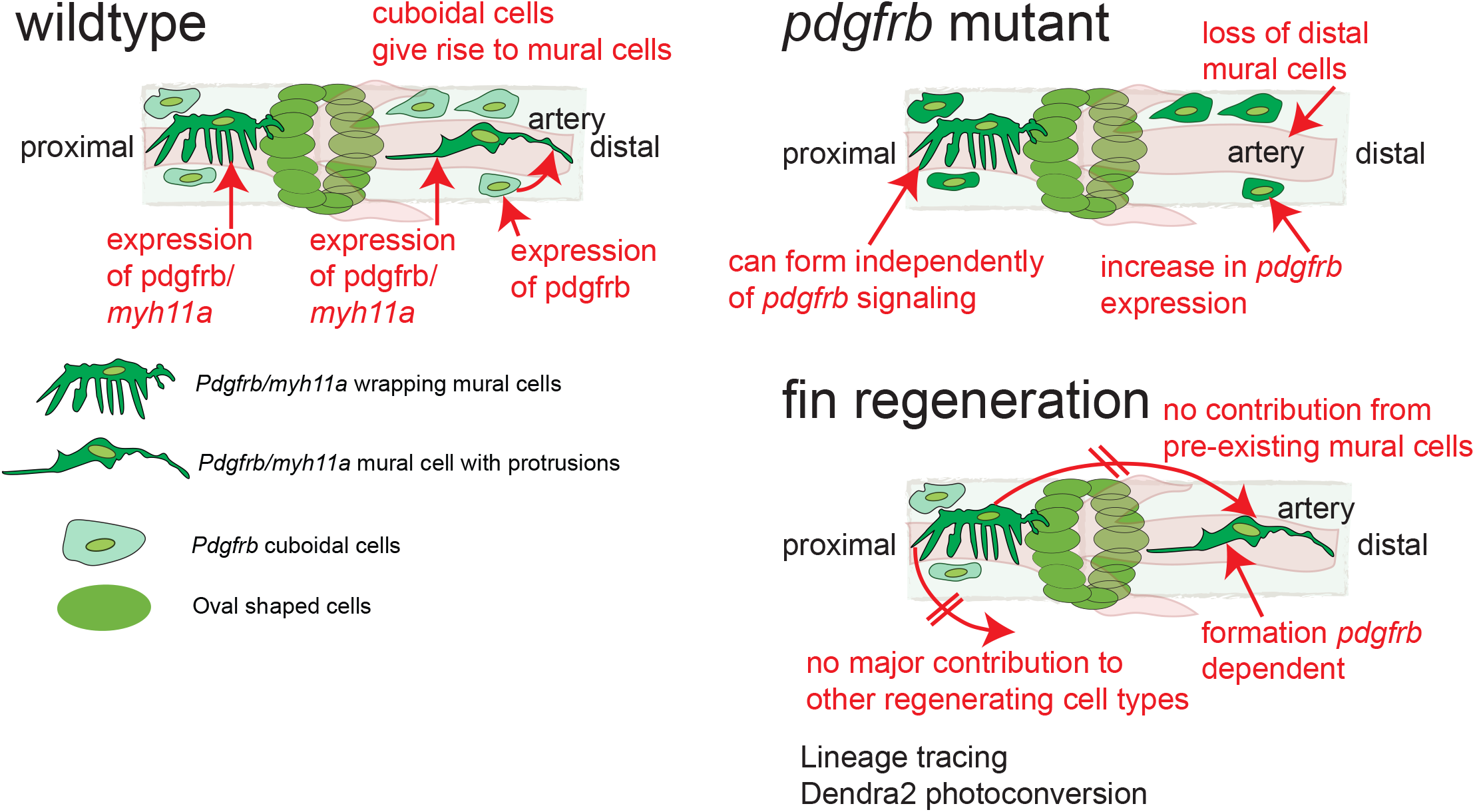
Graphical overview of findings.

